# Control with Practical Guarantees of Stationary Variance in Stochastic Chemical Reaction Networks

**DOI:** 10.1101/2025.05.12.653489

**Authors:** Armin M. Zand, Ankit Gupta, Mustafa Khammash

## Abstract

Biomolecular integral feedback controllers offer precise regulation of molecular species copy numbers, making them valuable for synthetic biology applications. Antithetic integral feedback controllers, in particular, can be effective in low-copy-number regimes with stochastic dynamics. In this work, we introduce a modified variant of this controller, called the antithetic dual-rein integral feedback motif, and analyze its performance from a stochastic perspective in the presence of intrinsic dynamic randomness. We demonstrate that our controller enables first-moment control while maintaining a tractable steady-state variance bound under specific parametric regimes. Notably, this variance bound is tunable, as it depends solely on the controller parameters. We derive these results using stochastic model-order reduction and validate them through numerical simulations. Our findings provide new insights into achieving both precise regulation and noise suppression in stochastic genetic circuits.

## I. Introduction

Synthetic biology is transforming the way we understand and engineer living systems by uniting molecular biology with concepts from control theory, computer science, and systems engineering. This interdisciplinary approach has enabled the rational design of synthetic genetic circuits that program cells to sense their environment, compute logical operations, and execute coordinated responses [1], [2]. Achieving robust control of cellular processes remains fundamentally challenging due to the inherent complexity and stochasticity of biological networks. Nevertheless, it holds immense promise for advancing synthetic biology, enabling the design of reliable and predictable biological systems (see [3]–[6] and references therein).

Chemical reaction networks (CRNs) serve as a powerful abstraction for modeling genetic circuits and biochemical interactions and have been successfully used to apply engineering principles and tools to cellular systems. The analysis of CRNs can be approached through two complementary perspectives. In the deterministic setting, the assumption of large molecular copy numbers enables the use of kinetic equations and differential equations to describe the network dynamics, where fluctuations due to molecular noise are negligible. However, biomolecules in single cells often exist in low copy numbers, necessitating a stochastic description to account for intrinsic noise [7]. The study of stochastic CRNs may reveal unique dynamical behaviors that are absent in deterministic settings, and vice versa [8]. Investigations into how feedback control strategies can be realized within the CRN framework has yielded a wealth of theoretical insights and experimental implementations [9]–[24].

Integral feedback controllers have been widely studied in this context, offering a mechanism for achieving disturbance rejection, a property commonly referred to as robust perfect adaptation (RPA) in biological literature, in genetic circuits [25]. Various such motifs have been characterized and studied in the literature, namely those relying on autocatalysis [24] or sequestration reactions (see [20] and references therein). Among these, the antithetic integral feedback motif was theoretically shown to be a necessary architectural component for achieving RPA at the mean level in stochastic regimes [12], [16]. This motif has been extensively analyzed in both deterministic and stochastic settings in terms of stability, performance, and output variance [26]–[39], leading to new insights as well as several different variants and extended designs.

While the classical antithetic motif can provide effective regulation, its performance in deterministic regimes can be compromised by uncertainties affecting the process, and in stochastic regimes, effective mean-level regulation of the output variable may come at the cost of large variances. To address these key limitations of its predecessor, we introduce a variant of the antithetic controller, which we name “the antithetic dual-rein integral feedback motif”. The present contribution aims to tackle the issue of large (unregulated) variance in the stochastic setting, while a complete analysis of the antithetic dual-rein control in the deterministic setting—including robustness results and performance guaran-tees—will be presented elsewhere.

Through stochastic analysis of the antithetic dual-rein motif and by exploring its moment dynamics (its mean and variance dynamics), here we show that this multi-variable biocontrol mechanism can be harnessed to drive the expected copy number of an output of interest to a predetermined, tunable set-point while ensuring practical guarantees of control over the copy number variance. By *practical* guarantees of variance, we mean that the controller provides a means of controlling the stationary variance under certain asymptotic limits and ensures it is always below certain (tunable) threshold, independent of the process reaction network. We will show that, under specific parameter regimes, our controller is able to achieve the first-moment regulation while maintaining an upper bound on the output variance, where this upper bound is also tunable solely based on the design (controller) parameters. To the best of our knowledge, this is the first result providing such theoretical guarantees on both mean and variance control, which may open avenues for exploiting biomolecular controllers in robust and precise control applications. The main results and conclusions are presented after we review some preliminaries.

## II. Preliminaries

### A. Chemical Reaction Networks

Here, we briefly review the framework of CRNs. Consider a CRN consisting of *l* different species **X**_**1**_, **· · ·**, **X**_**L**_ that react with each other through *m* distinct reaction channels ℛ_*j*_, which we specify in the form:

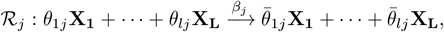

where *θ*_*ij*_ and 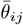 represent the molecular counts of reactants and products involved in the reaction and are therefore non-negative integers. In the deterministic setting, we associate a dynamical system with this reaction network, where the state variables *x*(*t*) = [*x*_*i*_(*t*)] ∈ ℝ^*L*^, *i* ∈ {1, · · ·, *L*} represent the species’ concentrations and evolve according to the dynamic model: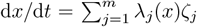, where 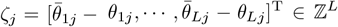 is the *stoichiometric vector* and *λ*_*j*_(*x*) is the rate or the *propensity* function whose choice depends on the kinetics used to model reaction *j*. For mass-action kinetics, for example, it is given by 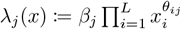.

In the stochastic setting, the state *x*(*t*) = [*x*_*i*_(*t*)] is a vector of non-negative integers, where *x*_*i*_(*t*) represents the copy number of species **X**_**i**_ at time *t*. Moreover the propensity function *λ*_*j*_(*x*) determines the rate at which reaction ℛ_*j*_ occurs and displaces the state by the stoichiometric vector *ζ*_*j*_. We model the underlying dynamics using a continuous-time Markov chain (CTMC), where the copy number vector *x*(*t*) evolves through time according to the transition rate matrix set by the propensity functions *λ*_1_, …, *λ*_*m*_ [40]. For mass-action kinetics, these propensity functions have the form 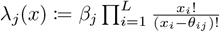.

### B. Considered Closed-loop Control Circuit

Here, we introduce the antithetic dual-rein integral feedback motif. This motif builds upon the original antithetic motif for the computation of the integral error but includes two opposing key actuation reactions that act on the output of interest (potentially indirectly through intermediate pathways), which we will later exploit to establish our theoretical guarantees. A poorly known CRN made of *n* different species, called X (t), is to be controlled by this controller. This CRN, representing the process network under control, may be subject to uncertainties and exogenous disturbances. It has two species through which it can communicate with the controller: **X**_**1**_ and **X**_**n**_. The latter species **X**_**n**_ is the species of interest, also referred to as the output species, whose quantity the controller aims to regulate. Under standard assumptions of set-point admissibility and stability, the output variable is steered to the value *µ/ρ* despite persistent disturbances (in the stochastic settings, this holds at the mean level).

The two actuation reactions that form the core components of this multi-variable biocontrol mechanism are indicated by red arrows in Fig. 1, and we shall write them as

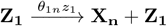

and

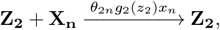

where the quantity above the arrow indicates the reaction rate and *g*_2_ is a design function that needs to satisfy certain conditions that we specify later. We shall define a small parameter called *ϵ* and rewrite the above two rate constants *θ*_1*n*_ and *θ*_2*n*_ based on it as

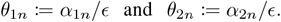

**Fig. 1.**
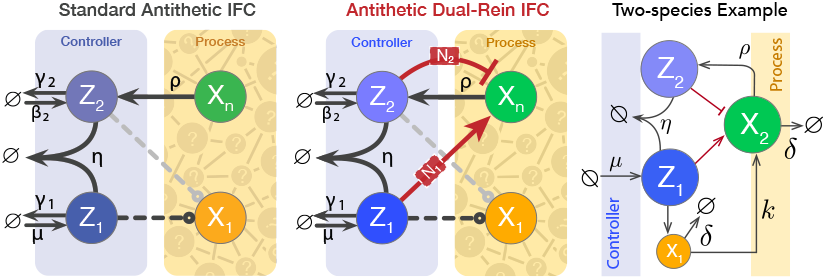
Antithetic dual-rein integral feedback controller motif. Motif depiction from left to right: the standard antithetic control, antithetic dual-rein control, and a gene-expression control example circuit.

The controller senses the output of the interest by the sensing reaction

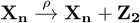

and computes the integral action by processing on the feed-back error through the following sequestration and set-point encoding inflow reactions

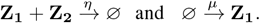

Throughout, we shall assume that the effects of controller species’ degradation/dilution due to cell growth and division are negligible, as are their basal expression rates (the rates *γ*_*i*_ and *β*_*i*_ in Fig. 1 are zero).

### C. Problem Statement

In this work, we focus on the stochastic analysis of the introduced controller motif by modeling the reaction network under consideration as a continuous-time Markov chain. Our goal is to derive analytical expressions for the stationary variance of the regulated copy number and to investigate whether theoretical guarantees regarding the control performance of our antithetic dual-rein controller can be established in the stochastic setting. Ergodicity of the closed-loop Markov chain, the well-posedness of the moment equations (i.e., finite variance), and set-point admissibility—that is, the controller’s ability to steer the expected copy number of the output variable of interest to the considered set-point under a stationary distribution—are our assumptions throughout. In all our simulation results hereafter, we take the process to be a two-species activation cascade (also referred to as the gene-expression example), where the expression of the first species (an mRNA) is induced by the actuation reaction from **Z**_**1**_, and this species, in turn, induces the expression of the output of interest (a protein). The closed-loop control circuit is comprised of four species and *n* = 2.

See Fig. 1 for graphical depiction. Let us write the rate at which the direct actuation reaction from **Z**_**1**_ on **X**_**1**_ takes place by *θ*_11_. The dual-rein feedback mechanism acts on the output species through both controller species. We limit our consideration to cases where these two actuation reactions are direct, meaning no intermediary species mediate the signals, and without loss of generality we allow further actuation reactions from the controller side on the input species **X**_**1**_ as well. Note, however, that similar results are expected to hold in indirect actuation cases, provided the intermediates within the networks N_1_ and N_2_ in Fig. 1 are at quasi-steady states.

For the numerical values used in our simulations, unless otherwise stated, we use the following default values: *ϵ* = 0.1, *η* = 10, *µ* = 9, *ρ* = 3, *θ*_11_ = 6, *α*_1*n*_ = 1, *α*_2*n*_ = 30 for the controller parameters; and *k* = 0.5 and *δ* = 2 for the process parameters. The set-point around which we expect the controller to regulate the mean of the output random variable is determined by the ratio *µ/ρ*, and its default value is therefore set to 3. For the function *g*_2_, we choose a monotonically increasing form as

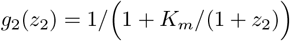

which could be realized, for example, if **Z**_**2**_ inhibits an intermediate complex that inhibits the expression of **X**_**n**_. To obtain our stochastic simulation results, we use the Gillespie algorithm (also known as the stochastic simulation algorithm or SSA), a widely used Monte Carlo-based approach for analyzing CRNs [41]. As for the sample size, we generate 10000 trajectories per simulation.

## III. Results: Stochastic Analysis of the Antithetic Dual-Rein Integral Controller Motif

In this section we provide the analysis of molecular interactions realizing the controller motif discussed in Section II-B, with now a stochastic perspective treating the whole system as a continuous-time Markov chain defined on some probability space (Ω, ℱ, ℙ), where Ω, ℱ, and ℙ are the sample space, associated *σ*-algebra, and the probability measure, respectively. In this paradigm, the random state variables of the Markov chain represent network species’ copy numbers, under well-mixed environment assumptions [40]. We shall assume massaction kinetics throughout. Let us use the capital letter *S*_*i*_ to associate a random variable to the species **S**_**i**_, denoting its copy number. We will accordingly use lower letter *s*_*i*_ to denote a particular realization of this random variable.

### A. Stochastic Model-order Reduction Analysis of the Closed-loop Control System

The states of the full system will evolve on a (*n* + 2)- dimensional non-negative lattice. The infinitesimal generator of this Markov chain can be written out as follows

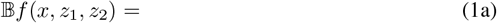

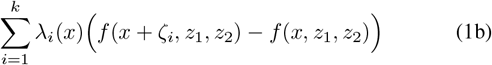

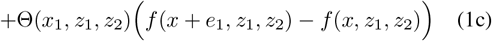

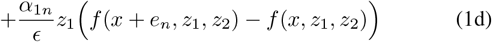

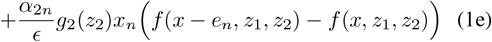

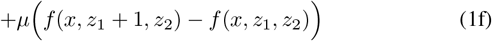

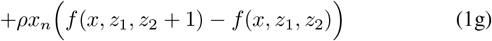

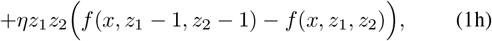

where *λ*_*i*_ represents the propensity function for the *i*-th reaction of the controlled process, ***X*** (*t*). The reactions within the network ***X*** (*t*), that are, the open-loop reactions, consist of *k* different reaction channels in total, 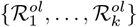 We assume no knowledge of the exact descriptions for the propensity functions *λ*_*i*_s and stoichiometric vectors *ζ*_*i*_s, which denote the copy number change upon firing the corresponding reaction 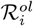. The propensity function Θ represents the effect of other (free) actuation terms on the species **X**_**1**_. We modeled it as a single production reaction (the vector *e*_*i*_ represents the standard basis vector with its *i*th element equal to unity), but it is fairly straightforward to reformulate the above generator for various other possible scenarios. In the gene-expression example case that we consider in this paper, it takes the form Θ(*x*_1_, *z*_1_, *z*_2_) = *θ*_11_*z*_1_.

The operator 𝔹*f* (*x, z*_1_, *z*_2_) above captures the infinitesimal changes of the distribution of the state random variables of the Markov chain over time, for any given real-valued function *f*. At stationarity, assuming the ergodicity of the network (or other notions of stochastic stability) with a unique stationary distribution *π*(*x, z*_1_, *z*_2_), the following holds

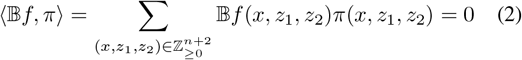

for a class of polynomially growing [42] real-valued functions *f*. Our ideal goal, though may not be easily achievable due to the nonlinear propensities and the ensuing moment closure problem [43], is to find the analytical stationary distributions (at least up to the second moment) of the regulated jump random variable *X*_*n*_. For the first moment, it is easy to observe from the dynamics that 𝔼_*π*_(*X*_*n*_) = *µ/ρ*, under our assumption of ergodicity of the closed-loop network.

Let us first derive a valid quasi-stationary model in the case *ϵ* → 0 for the system described by (1) under suitable assumptions. As *ϵ* gets smaller, the full system described by (1) will have two time scales for the reaction dynamics. The fastest time scale will be governed by two reactions corresponding to the terms (1d) and (1e) in the generator. These two fast reactions only affect the output state *X*_*n*_ according to the birth-death process:

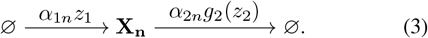

The rest of the reactions will occur at the slow time scale, and in the limit ∈ → 0, we can make the standard quasi-stationarity assumption where all the slow variables (i.e. all state variables except *X*_*n*_) will be kept constant. To have the guaranteed convergence (in distribution) of *X*_*n*_ from the full system in (1) in the limit *ϵ* → 0 to that of obtained from the birth-death process (3), we would need to meet some technical assumptions for (3), including that it remains ergodic for every choice of the (*z*_1_, *z*_2_) pair, according to [44]. Different other works have also been looking into this stochastic quasi-stationarity approximation problem, from different points of views [45], [46]. We shall take the following as given throughout this section, which is a boundedness type of assumption on *g*_2_. Note that, for the quasi-stationarity analysis of this section to be valid, we do not make any other assumptions about the function *g*_2_, including those of monotonicity.

#### Assumption 1.

*There exists a positive constant* 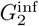 *such that* 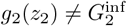 *for all z*_2_ ∈ ℤ_≥0_.

It is easy to check that the cases where *g*_2_ satisfies Assumption 1, it ensures that the quasi-stationarity assumption with the fast dynamics for *X*_*n*_ defined by (3) is valid. See [44] and Remark 4.5 therein. This way we arrive at a reduced Markov process, that does not involve the output species state, and can approximately capture the probability evolution of the full system (1) over compact time intervals in the limit ϵ → 0. The generator of this reduced system may be written as follows

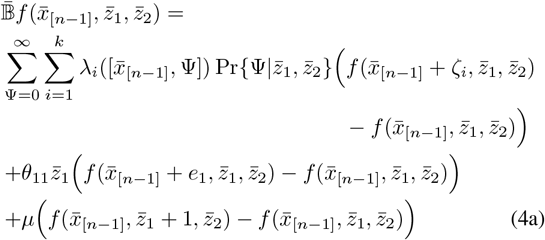

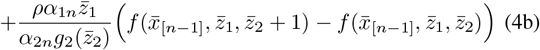

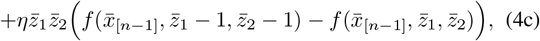

where Ψ is an auxiliary discrete random variable whose probability mass function 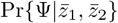 is given by

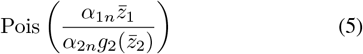

where Pois(**•**) represents the distribution parameterized with **•**. We use the over-bar symbol to distinguish the corresponding random variables from the reduced system. Here, 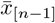 represents the reduced process state vector 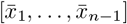. The random variable Ψ is an approximation of *X*_*n*_ from the full system for small ϵs, and we will always compare its distribution with that of *X*_*n*_ over time. While the model-order reduction results are rigorously valid over compact time intervals, we shall assume throughout this study that in the ergodic limit as *t* → ∞ it remains valid too, over at least an interval of ϵ ∈ (0, *ϵ*_*d*_), with *ϵ*_*d*_ ∈ ℝ_*>*0_ a constant. Notice that the state space lattice on which the jump process in (4) evolves has been reduced to 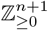. Notice also that the term 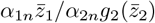 in (4b) appears as the solution of 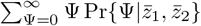, which admits this simple closed-form expression because the corresponding propensity function is linear and the distribution of Ψ is Poisson. The simulation results provided in Fig. 2, Fig. 3, and Fig. 4 verify the findings of this subsection and demonstrate the closeness of the full and reduced systems as *ϵ* approaches zero. In the next subsection, we focus on the reduced Markov process (4) and find analytical bounds on its stationary variance by closing the moments under specific parametric regimes.

**Fig. 2.**
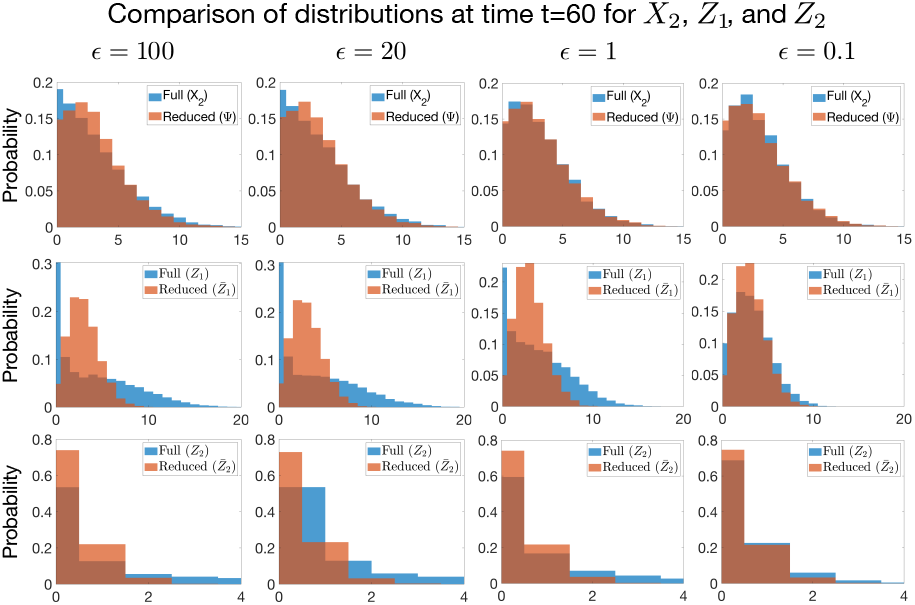
The distribution of the full system’s copy numbers converges to that of the reduced system as *ϵ* approaches zero. For four different values of *ϵ*, the histogram plots at each row compare the distributions of the copy numbers *X*_2_, *Z*_1_, and *Z*_2_ at a single time point with the corresponding distributions obtained from the reduced system, namely those of the random variables Ψ, 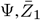, and 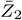, respectively. *K*_*m*_ was set to 0.001 and *α*_2*n*_ = 1.

**Fig. 3.**
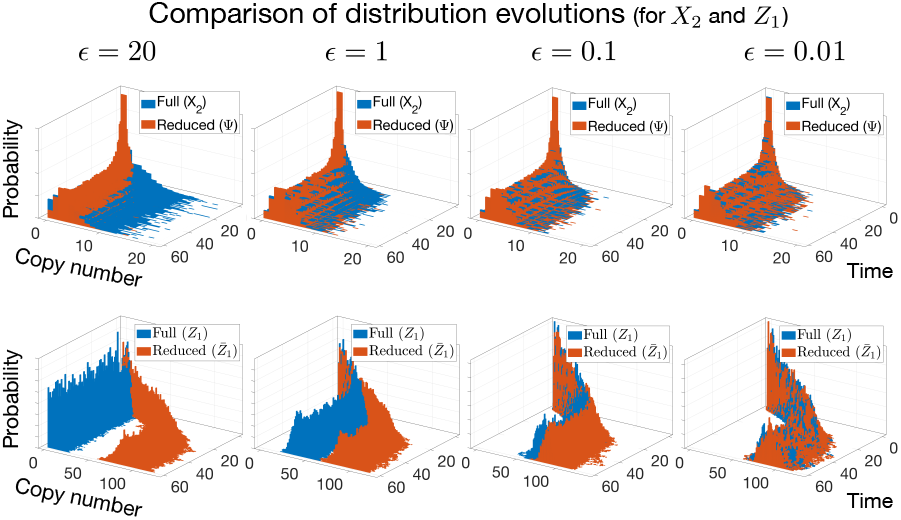
Full time-interval distribution comparison corresponding to the distributions and simulation results in Fig. 2, but for a different value of *α*_2*n*_ (*α*_2*n*_ = 30) and smaller *ϵ*s. Instead of a single discrete time point, bivariate histograms are used to provide a full time-spectrum comparison (*t* ∈ [0, 60]). Each *z*-*y* slice of the 3D plot at *x* = *t* illustrates the probability distribution of the corresponding copy number at time *t*.

**Fig. 4.**
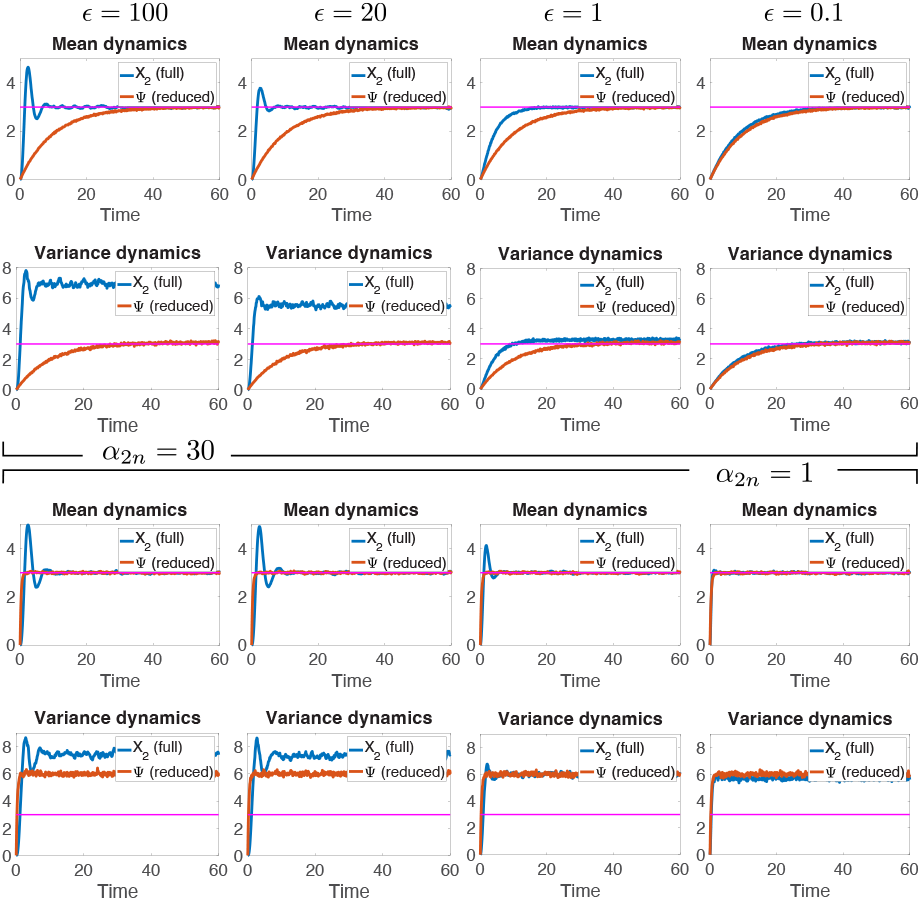
The mean and variance dynamics of the output random variable from the full system compared to those of the random variable Ψ. The upper panel corresponds to the system from Fig. 3, while the lower panel presents the same simulations for a different value of *α*_2*n*_. The reference line in pink represents the set-point value *µ/ρ* (which is equal to 3), providing a comparison with the Poissonian-level noise at stationarity.

### B. Analytical Approximation of the Stationary Variance in Certain Cases

In this subsection, we work with the reduced generator 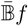 in (4) and try to approximate its stationary distribution. We work with a general process ***X*** (*t*) and take the controller design parameters as some fixed, given values, except the ratio between the two tuning gains *α*_1*n*_ and *α*_2*n*_. First, let us impose a stricter form on the function *g*_2_ than the one presented in Assumption 1, that is, we allow *g*_2_s that are both upper and lower bounded by some positive, known constants.

#### Assumption 2.

*The function g*_2_(*z*_2_) *is chosen such that* 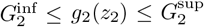 *for all z*_2_ ∈ ℤ_≥0_, *where* 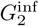 *and* 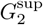 *both are some positive fixed constants*.

Note that, from the form of the generator in (4), the controller-related propensity functions have now been decoupled from the process random variables and do not depend on 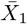 to 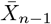, but only depend on the two random variables 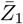 and 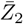. This reduces the problem at hand to only considering the evolution of the distributions in the two-dimensional 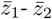 lattice in order to gain enough information regarding the moment distributions of the random variable of interest; here Ψ. Yet it is challenging to close the moments for such a decoupled, lower-dimensional subsystem due to the presence of nonlinear propensities, as can be seen from (4). We would thus need to tackle the moment closure problem by considering some specific parametric regimes.

Before continuing, let us provide some properties of the invariant distributions for the decoupled two-dimensional sub-system. By considering to solve from (2) for different choices of functions 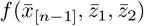 according to 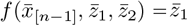 and 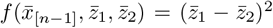, one can, respectively, obtain the next set of equalities at stationarity

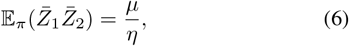

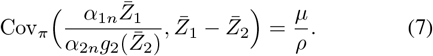

The proof of the above result has been omitted because it follows a similar procedure as in [31, Supplementary Material]. Under the assumption that the closed-loop network is ergodic (convergence to a unique distribution in the limit *t* → ∞), the controller will ensure that at steady-state the following expression holds

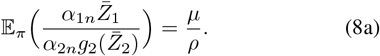

From the law of total expectation, this follows that 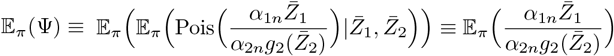 and hence

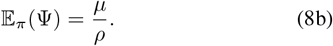

This means that the first-moment regulation that holds for the full random variable *X*_*n*_ under ergodicity and set-point admissibility assumptions also holds for its reduced-model approximation Ψ, thanks to the fact that the structure of a functional integrator is preserved in the reduced stochastic system after model-order reduction is applied. Next, we exploit the law of total variance to prove the following lemma.

#### Lemma 1.

*Given Assumption 1, the stationary variance of the random variable* Ψ *is bounded from above according to the stationary variance of* 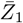 *such that*

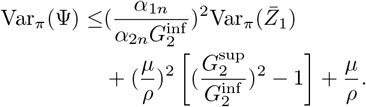

*Proof*. We know that 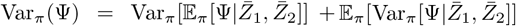. This equivalently follows

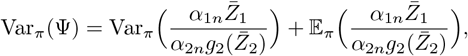

given the fact that Ψ’s distribution follows a Poisson distribution parameterized according to (5). The latter term is equivalent to *µ/ρ*, due to (8). Now, we are left to upper bound the first term 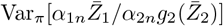 based on 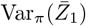. From Assumption 1, we have that for any pair 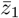 and 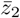, the value of 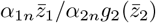 is always less than or equal to 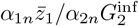 and greater than or equal to 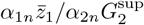. This means that the first and second moments of the random variable 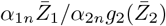 are upper bounded by those of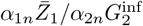. Moreover,

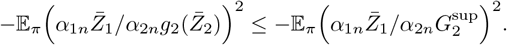

Thus,

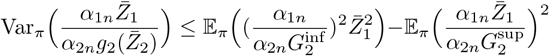

which is less than or equal to 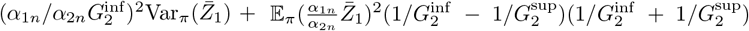. From (8), we know that at stationarity 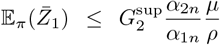 and thus 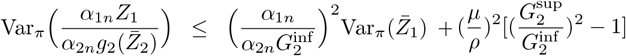.

Also, we consider the following result, which is valid in the limit of small *α*_1*n*_*/α*_2*n*_ ratio.

#### Lemma 2.

*Under Assumption 2 and for sufficiently large but finite values of the ratio α*_2*n*_*/α*_1*n*_, *we have the following valid at stationarity for the random variable* 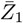

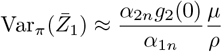

*Proof*. From Assumption 2, we know that the random variable 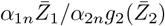 satisfies

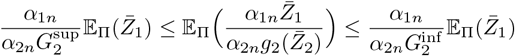

at any distribution Π. From the equation (8), we get at stationarity that

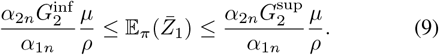

This means, for sufficiently large but finite values of the ratio *α*_2*n*_*/α*_1*n*_, the expected value of 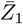 has to take large values. See Fig. 5 for simulation results verifying this. Using this fact, we refer to the invariant equation (6) as to express a possible way closing the moment equations. From the stationary equation wherein, it is immediate that, as 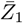 takes larger values on expectation, 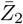 has to be having values closer to 0 at stationarity, in order that the equality in (6) is met (assuming that we only change *α*_1*n*_ and *α*_2*n*_, nor any of *µ* or *η*). See Fig. 5 for numerical verification. Consequently, we can write 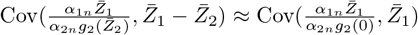. By referring to (7), one can write the approximate expression

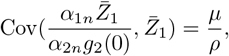

which equivalently means 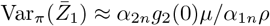.

**Fig. 5.**
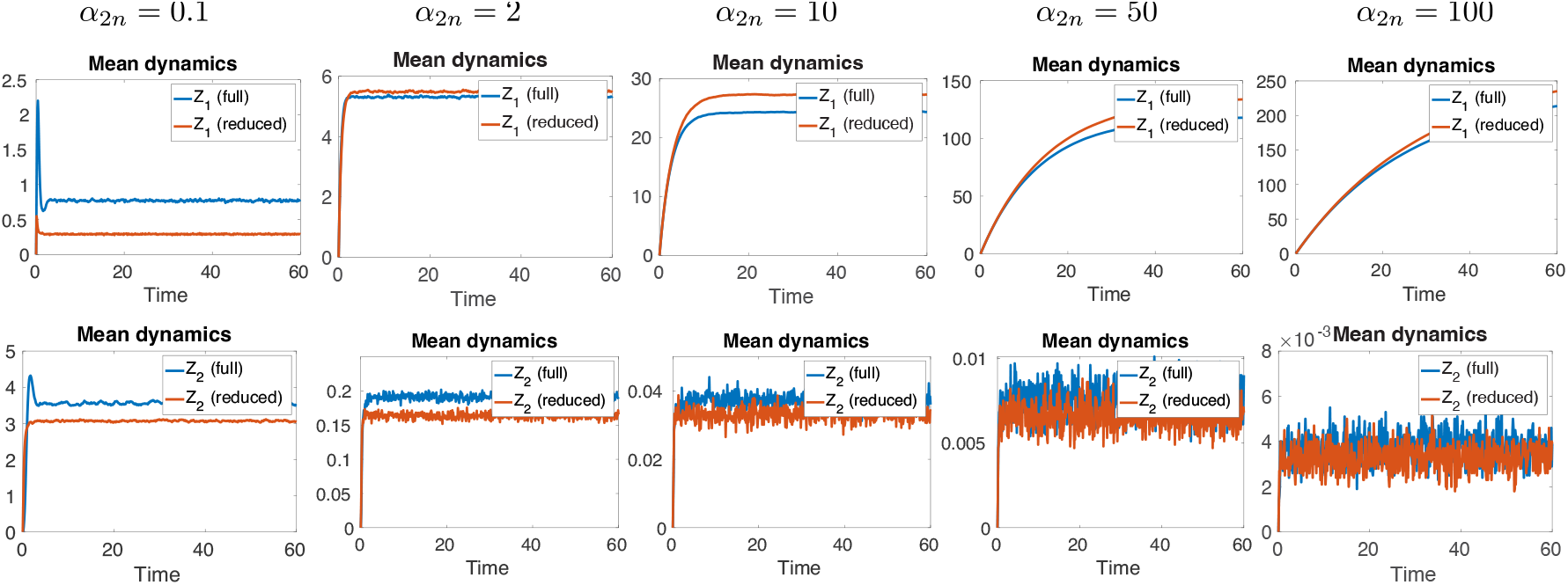
Top row: The expected value of 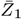, and consequently *Z*_1_ for sufficiently small *ϵ*, increases as *α*_2*n*_ increases. Bottom row: The random variable 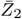, and consequently *Z*_2_ for sufficiently small *ϵ*, takes values closer to zero as *α*_2*n*_ increases. This approximation is used in the proof of Lemma 2 as part of a moment-closure approach. The parameter *K*_*m*_ was set to 0.1.

Now, we are in a position to state the main result of this subsection.

#### Proposition 1.

*Suppose Assumption 2 holds. For sufficiently large but finite values of the ratio α*_2*n*_*/α*_1*n*_, *the approximate stationary variance of the random variable* Ψ *satisfies*

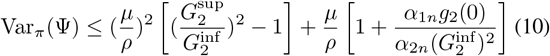

*Proof*. The proof easily follows if taken together the statements of Lemma 1 and Lemma 2.

The above result provides an explicit upper bound on the stationary variance of the regulated random variable, which is tunable solely based on the controller (design) parameters. Interestingly, this upper bound approaches arbitrarily close to the mean value as *K*_*m*_ tends to zero and the ratio *α*_2*n*_*/α*_1*n*_ tends to ∞ (given the chosen function for *g*_2_, which represents the inhibition of **X**_**n**_’s inhibition by **Z**_**2**_). Recall that we chose *g*_2_(*z*_2_) = 1*/*(1 + *K*_*m*_*/*(1 + *z*_2_)), where *K*_*m*_ is a parameter associated with this function (here, 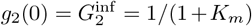 and 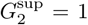). We expect this stationary variance bound to hold analogously for the full system’s output variable *X*_*n*_ when *ϵ* is sufficiently small. Fig. 6 includes simulations verifying these findings, wherein the green lines are used to indicate the corresponding upper bound calculated from (10).

**Fig. 6.**
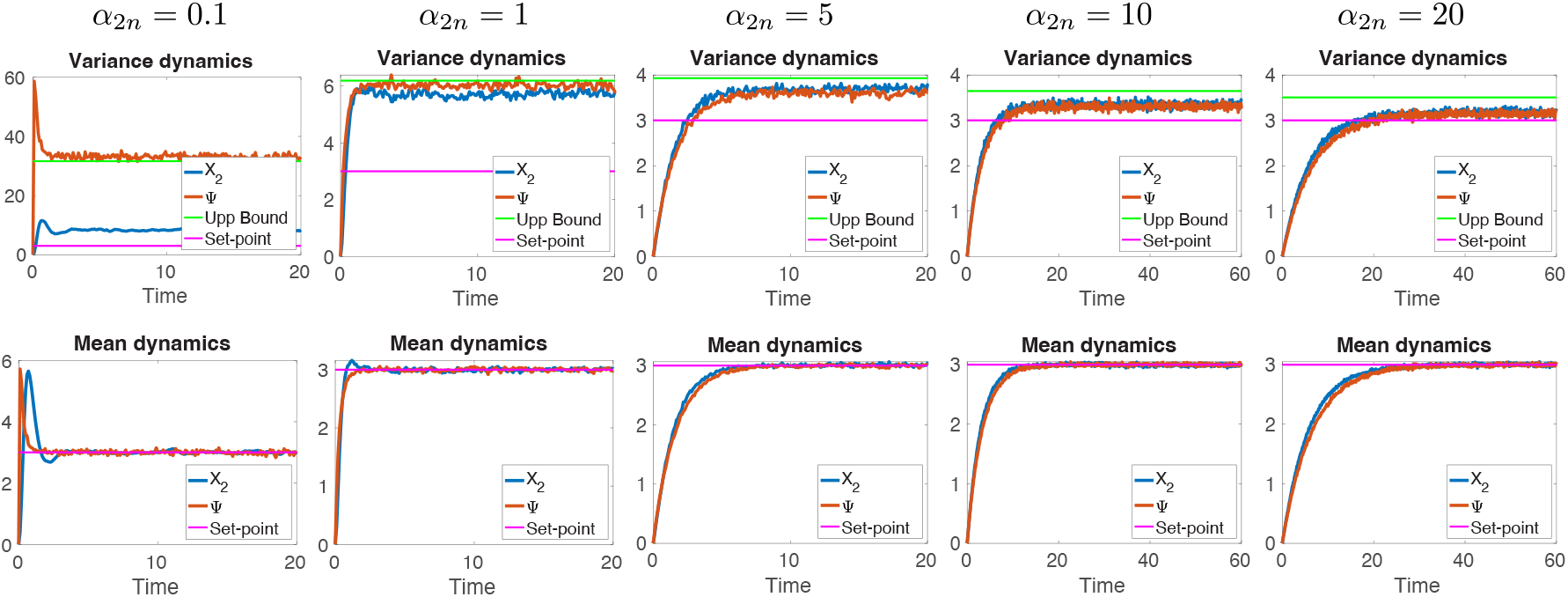
By choosing a sufficiently large *α*_2*n*_*/α*_1*n*_ ratio and a sufficiently small *ϵ*, the antithetic dual-rein feedback controller ensures that the stationary variance of the output copy number remains below a fixed, tunable upper bound (shown in green and derived from the right-hand side formula in (10)), while tightly regulating its stationary expected value about the set-point *µ/ρ*. The parameter *K*_*m*_ was set to 0.02.

## IV. Conclusion

In this paper, we introduced the antithetic dual-rein integral feedback motif and analyzed its performance in the presence of dynamic randomness and intrinsic noise. Here, we investigated whether certain theoretical guarantees hold for the moments of the output variable in stochastic settings. To this end, we derived a reduced-order model under the assumption of a small parameter ϵ approaching zero. The resulting boundary-layer solution indicated that the stationary distribution of the output random variable converges to a Poisson distribution, denoted by Ψ, whose parameter depends on two random variables with distributions dynamically evolving over time. We imposed boundedness assumptions on the function *g*_2_(*z*_2_) to support our derivations.

At the mean level, and under the assumption of ergodicity, the considered controller steers the expected value of the output copy number to the set-point value *µ/ρ*. Additionally, we derived an upper bound on the (approximate) stationary variance of the reduced output Ψ in terms of the controller parameters. We developed a moment-closure method to approximate the first two stationary moments of Ψ in certain parametric limits, and simulation results supported the validity of the obtained variance bound. For sufficiently small *ϵ*, the variance of the full output variable closely approximates that of the reduced one, thereby remaining below the obtained upper bound.

These findings highlight the ability of the introduced antithetic dual-rein control strategy to manage both mean-level regulation and control over variability (arising from intrinsic noise) in a fully tunable manner, independent of the disturbances or uncertainties affecting the controlled process. We also discussed specific parametric tuning strategies under which the obtained upper bound approaches the mean level, implying near-Poissonian moment control in stochastic reaction networks. Numerical simulations on a two-species gene expression model validated our analytical derivations. However, our analysis relied on the assumption that the reduced system is ergodic and that its distribution closely approximates that of the full system at all times. A mathematically rigorous proof of the validity of the model-order reduction and the ergodicity of the reduced system requires deeper theoretical investigation and is left for future work. Such further analysis and extended results would help characterize the precise parametric and structural properties of process networks under which the obtained variance bound remains valid at stationarity.

## V. Acknowledgments

This work was supported by the Swiss National Science Foundation (SNSF) Advanced Grant: Theory and Design of Advanced Genetically Engineered Control Systems (grant number 216505). The authors used ChatGPT (by OpenAI) to improve the language and readability of the manuscript. They take full responsibility for the content and its interpretation.

## Notes

### Competing Interest Statement

The authors have declared no competing interest.

